# Neurodegeneration induces a developmental RNA processing program by calpain-mediated MBNL2 degradation

**DOI:** 10.1101/2020.07.24.219121

**Authors:** Lee-Hsin Wang, Yu-Mei Lin, Chien-Yu Lin, Yijuang Chern, Guey-Shin Wang

## Abstract

The Muscleblind-like (MBNL) protein family plays an important role in regulating developmental RNA processing transition. Loss of MBNL2 function has been implicated in the neurodegeneration of myotonic dystrophy type 1 (DM1). However, the causal mechanism of neurodegeneration-induced MBNL2 loss of function remains elusive. Here, we show that neurodegenerative conditions including NMDAR-mediated excitotoxicity and dysregulated calcium homeostasis triggered nuclear translocation of calpain-2 resulting in MBNL2 degradation and reversion of MBNL2-regulated RNA processing to developmental patterns. The developmental stage featured nucleus-enriched distribution of calpain-2 and low expression of MBNL2. Increased MBNL2 expression during development is required for promoting developmental RNA processing transition and neuronal maturation. Knockdown of calpain-2 expression inhibited neurodegeneration-induced MBNL2 reduction and dysregulated RNA processing. Neurodegenerative disease mouse models including DM1 and Alzheimer’s disease showed nuclear translocation of calpain-2 associated with MBNL2 degradation and reversion of MBNL2-regulated RNA processing to the developmental pattern. Our results identify a novel regulatory mechanism for MBNL2 downregulation and suggest that reduced MBNL2 expression accompanied by the re-induction of a developmental RNA processing program may be a common feature of neurodegeneration.

## Introduction

Regulation of gene expression at the post-transcriptional level, including 5’- and 3’-end processing, splicing, RNA editing and modifications, confers proper temporal and spatial gene expression as well as maintenance of physiological functions (1). Aberrant RNA processing, such as misregulated alternative splicing and polyadenylation, has been associated with neurological disorders including amyotrophic lateral sclerosis (2, 3), Parkinson’s disease (4), frontotemporal dementia (5), sporadic Alzheimer’s disease (6), Huntington’s disease (7) and myotonic dystrophy (DM) (8, 9). RNA-binding proteins (RBPs) play a critical role in regulating RNA processing events. In addition to mutations in RBP recognition sites, altered RNA processing under diseased conditions is mainly due to mutations in or dysregulation of RBPs (10, 11). Thus, understanding the causal mechanism of the perturbed function of individual RBPs in the diseased condition may help in developing a therapeutic strategy.

The Muscleblind-like (MBNL) protein family, including MBNL1, MBNL2 and MBNL3, plays an important role in mediating alternative splicing and polyadenylation, mRNA localization and miRNA processing (12-15). The MBNL protein family promotes differentiation by regulating the developmental alternative splicing transition (16). Loss of MBNL function has been implicated in the neural pathogenesis of DM type 1 (DM1) (14, 17). The genetic basis of DM1 is an expansion of CTG trinucleotide repeats in the 3’ untranslated region (UTR) of DM protein kinase (*DMPK*) gene (18). *DMPK* mRNA containing CUG repeats accumulates in the nuclear foci and binds and sequesters MBNL proteins, thereby resulting in loss of MBNL functions (8).

Despite the common feature of binding to expanded CUG RNA, members of the MBNL family exhibit different characteristics in response to expanded CUG RNA. MBNL1 exhibits the highest binding and mobility to the expanded CUG RNA and is required for RNA foci formation (19). In a brain-specific DM1 mouse model, EpA960/CaMKII-Cre, which expresses expanded CUG RNA in the postnatal brain, synaptic dysfunction associated with reduced cytoplasmic MBNL1 expression occurred before aberrant alternative splicing associated with the reduced MBNL2 expression (20). The finding suggested that expanded CUG RNA-induced misregulation of alternative splicing was likely due to reduced MBNL2 expression, which was consistent with findings from *Mbnl*-knockout mouse models showing that MBNL2 dysfunction played a major role in contributing to aberrant alternative splicing (14).

Reduced cytoplasmic MBNL1 is due to nuclear translocation resulting from deubiquitination of MBNL1 induced by expanded CUG RNA (21), but how expanded CUG RNA affects MBNL2 expression remains unclear. Detection of aberrant MBNL2-regulated splicing occurred after reduced integrity of axons and dendrites in EpA960/CaMKII-Cre mouse brains (20). Whether reduced MBNL2 expression was associated with or caused by neurodegeneration has not been determined. The major molecular characteristic of DM1 is misregulation of developmental alternative splicing and polyadenylation transitions (22); whether regulation of this transition is affected under neurodegeneration remains elusive. Moreover, whether misregulated RNA processing associated with reduced MBNL2 expression might be a common feature of neurodegenerative disorders has not been tested.

In the present study, we aimed to determine the regulatory mechanism of MBNL2 degradation and assess whether re-induction of the MBNL2-regulated developmental program could be a common feature of neurodegeneration. We first examined the temporal and spatial expression of MBNL2 during neuronal maturation and its involvement during neurite outgrowth by using cultured hippocampal neurons. Increased expression of MBNL2 during neuronal maturation was associated with the transition of MBNL2-regulated RNA processing toward the adult pattern. In two conditions revealing neurodegenerative features, glutamate-induced excitotoxicity and aberrant calcium homeostasis, induced nuclear translocation of calpain-2 degraded MBNL2 and induced reversion of MBNL2-regulated RNA processing into developmental patterns. During neuronal maturation, the distribution of calpain-2 followed a nuclear-to-cytoplasm transition, whereas expression of calpain-2 tagged with a nuclear localization signal promoted MBNL2 degradation. Thus calpain-2 nuclear translocation induced by neurodegenerative conditions resembled reversion to the developmental pattern. In addition, mouse models of neurodegeneration including DM1 and AD showed calpain-2 nuclear translocation and MBNL2 downregulation accompanied by dysregulated transition of developmental RNA processing. Depletion of endogenous calpain-2 upon neurodegenerative stimulation prevented MBNL2 degradation and reversion of RNA processing to the developmental stage. Our results suggest the re-induction of an MBNL2-regulated developmental program as a common feature of neurodegenerative disorders.

## Results

### MBNL2 is required for neuronal maturation and regulating the developmental transition of RNA processing

To determine the association of the MBNL2 expression pattern with neuronal maturation, we first examined MBNL2 expression during mouse brain development. MBNL2 expression was barely detected in early developing brains, including at embryonic day 14.5 (E14.5) and postnatal day 1 (P1) (Fig. 1*A*), but became detectable by P8 and increased afterwards during postnatal maturation (Fig. 1*A*, P15). The expression of MBNL2 was maintained throughout adulthood and was distributed in the cortex, hippocampus, striatum, amygdala and thalamus (Fig. 1*B*). Development of the granular neurons of dentate gyrus in the hippocampus follows an “ectal-to-endal” gradient pattern (23). We detected MBNL2 in the outermost granule cell layer of the external dentate limb (Fig. 1*C*, P8). As a function of time, the number of MBNL2-expressing cells increased in a gradient, from the external to internal layers of the dentate limb (Fig. 1*C*, P15), which indicates consistency with the pattern of maturation.

**Fig. 1.**
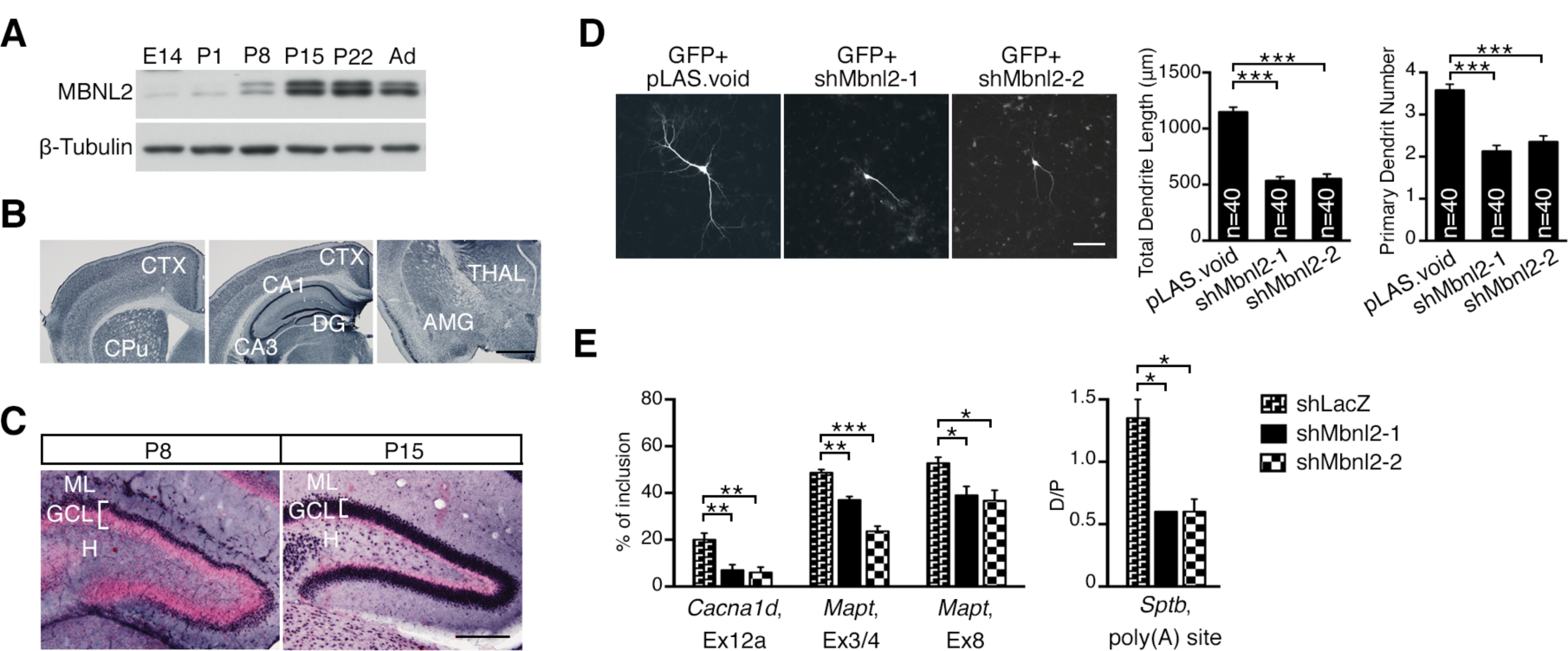
MBNL2 is required for regulating neuronal morphogenesis and the developmental transition of RNA processing. (*A*) Temporal expression of MBNL2 in the mouse forebrain at different ages including embryonic day 14.5 (E14.5), postnatal day 1, 8, 15, 22 (P1, P8, P15, P22) and 2 months (Ad). (*B*) MBNL2 immunoreactivity detected by immunohistochemistry in mice at age 4 months. (*C*) MBNL2 immunoreactivity with nuclear fast red used for nuclei labelling in the dentate gyrus of the P8 and P15 hippocampus. (*D*) MBNL2 knockdown impairs dendrite development. Cultured hippocampal neurons were transfected with plasmids expressing MBNL2 shRNA (shMbnl2-1 or -2) or control shRNA (pLAS.void). Plasmid expressing GFP was co-transfected to label neuronal morphology. Quantification of total dendrite length (left) and primary dendrite number (right). (*E*) Aberrant MBNL2-regulated RNA processing in Mbnl2-knockdown neurons. Quantification of the inclusion of *Cacna1d* exon 12a, *Mapt* exons 3/4 and 8, and the distal (D) to proximal (P) polyA utilization of *Sptb*. Data are mean ± SEM of indicated n number (*D*) or 3 independent experiments (*E*). \**P*<0.05, \*\**P*<0.01, \*\*\**P*<0.001, by one-way ANOVA (*D* and *E*). CTX, cortex; CPu, caudate putamen; DG, dentate gyrus; AMG, amygdala; THAL, thalamus; ML, molecular layer; GCL, granule cell layer; H, hilus. (Scale bar: *B*, 1 mm, *C*, 200 µm, D, 100 µm.)

We next determined whether MBNL2 plays a role in dendrite development. We first used two different *Mbnl2*-specific short hairpin RNA (shRNA) constructs to deplete MBNL2 in cultured hippocampal neurons. A GFP expression construct for showing the neuronal morphology was co-transfected with *Mbnl2*-shRNA constructs on day 11 of *in vitro* culture (DIV) and the dendrite development was analyzed at 14 DIV. MBNL2 knocked-down neurons exhibited shorter total dendrite length and reduced primary dendrite number as compared with control neurons, which suggests that loss of MBNL2 function may impair dendrite maturation (Fig. 1*D*).

We next determined whether MBNL2 depletion affected MBNL2-regulated RNA processing in cultured hippocampal neurons. During brain development and maturation of hippocampal neurons *in vitro*, the inclusion of *Cacna1d* exon 12a, *Mapt* exon 3/4 and exon 8 was increased, and utilization of *Sptb* alternative poly(A) sites was toward the distal site (*SI Appendix*, Fig. S1) (14, 20). However, in Mbnl2-depleted neurons, MBNL2-regulated RNA processing showed reversion to the developmental pattern; inclusion of the *Cacna1d* exon 12a, *Mapt* exon 3/4 and exon 8 was decreased, and utilization of *Sptb* alternative poly(A) sites was shifted toward the proximal site (Fig. 1*E*). Our results suggested that MBNL2 was required for dendrite development and regulation of RNA processing transition toward the adult stage.

### NMDA receptor-mediated excitotoxicity reduces MBNL2 expression

Reduced MBNL2 expression in a DM1, EpA960/CaMKII-Cre, mouse brain at 12 months of age is reminiscent of its pattern in an early developmental stage (20). The causal mechanism of how MBNL2 expression is reduced in the EpA960/CaMKII-Cre brain is still unclear. To investigate this, we first determined whether neurodegeneration results in MBNL2 degradation. Glutamate-induced excitotoxicity is implicated in neurodegeneration (24). We used cultured hippocampal neurons to study the effect of glutamate-induced excitotoxicity on MBNL2 expression. Mature neurons cultured at 20 DIV were treated with 20 μM glutamate and harvested at different times. Glutamate-induced excitotoxicity was revealed by condensed nuclei on DAPI staining (25) (*SI Appendix*, Fig. S2). Next, we determined whether glutamate stimulation affects MBNL2 expression. At 1 hr after glutamate stimulation, MBNL2 protein level was decreased (Fig. 2*A*). With longer exposure time, MBNL2 was detected at lower molecular weight, from 20 to 40 kD, which suggests degraded products. With increased glutamate treatment time, MBNL2 expression continued to decrease. On immunofluorescence staining under the same exposure time, the intensity of MBNL2 immunoreactivity was reduced in neurons stained with anti-NeuN antibody upon glutamate stimulation (Fig. 2*B*).

**Fig. 2.**
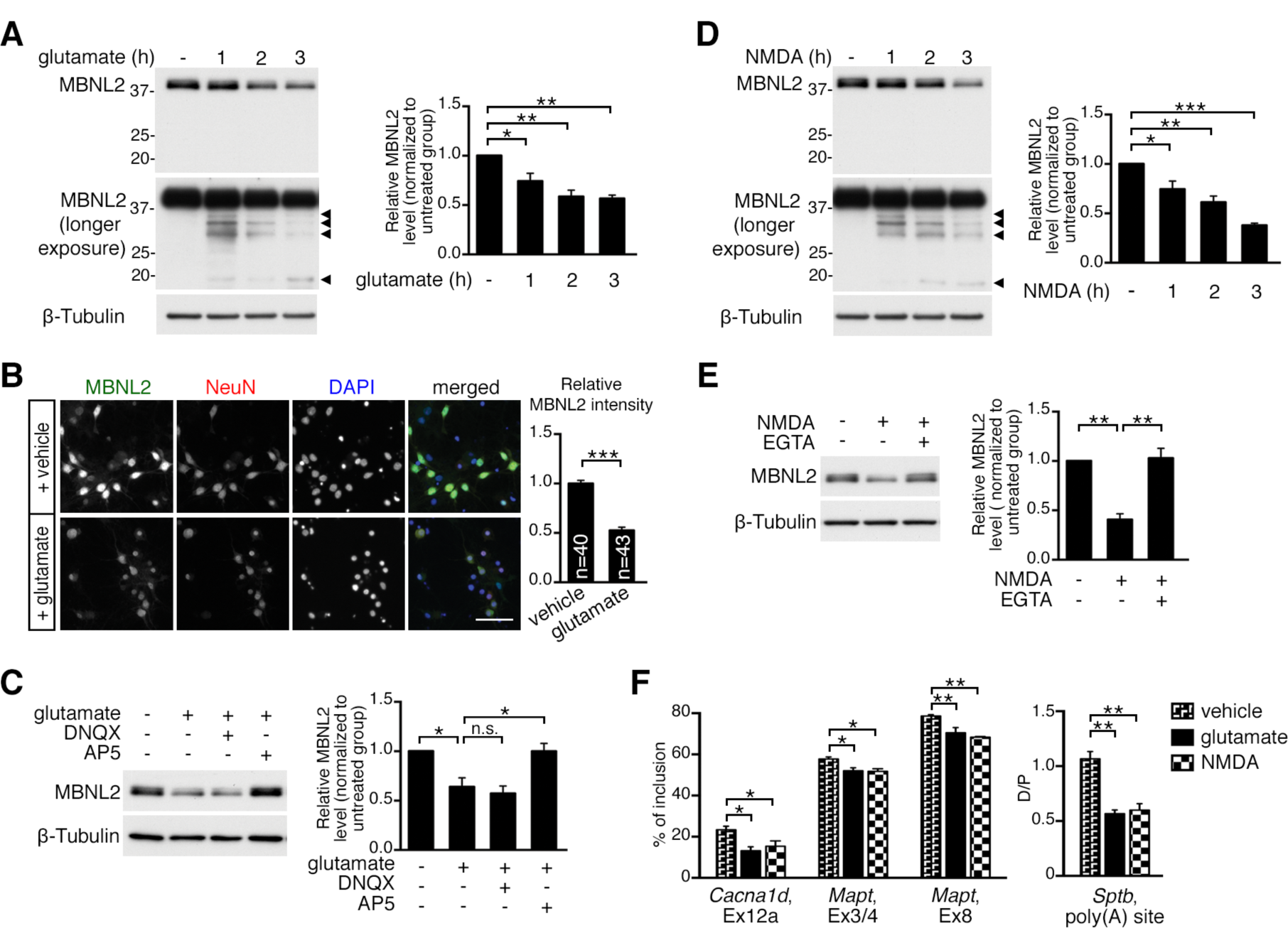
NMDAR-mediated excitotoxicity induces MBNL2 downregulation. (*A*) MBNL2 level was reduced in mature cultured hippocampal neurons treated with glutamate for the indicated times. MBNL2 level detected with longer exposure time is shown at the bottom. β-tubulin was a loading control. Relative amount of MBNL2 for treatments was compared by normalization with β-tubulin. Arrowheads indicate the possible degraded products of MBNL2. (*B*) Representative images of MBNL2 immunoreactivity in neurons treated with vehicle or glutamate. NeuN staining served as a neuronal marker. Quantification of overall MBNL2 intensity is shown at right. (Scale: 50 µm.) Number of neurons (n) used for quantification is indicated. (*C*) NMDAR antagonist AP5 but not AMPAR antagonist DNQX pretreatment preserved MBNL2 level in glutamate-treated neurons. (*D*) MBNL2 level was reduced in neurons treated with NMDA for indicated times. (*E*) Pretreatment with calcium chelator EGTA preserved MBNL2 level in NMDA-treated neurons. (F) Aberrant MBNL2-regulated RNA processing events in the glutamate or NMDA-treated neurons. Data are mean ± SEM of indicated n number (*B*) or 3 independent experiments (*A, C, D, E* and *F*). \**P*<0.05, \*\**P*<0.01, \*\*\**P*<0.001, by one-way ANOVA (*A, C, D, E* and *F*) or Student *t* test (*B*). n.s., not significant.

Glutamate-elicited excitotoxicity is mediated by calcium influx via the activation of NMDA receptor (NMDAR) (24). We next determined whether NMDAR activation is involved in glutamate-reduced MBNL2 expression by pretreating neurons with AP5 or DNQX, an NMDAR and non-NMDAR antagonist, respectively, before glutamate treatment. Pretreatment with AP5 but not DNQX blocked glutamate-reduced MBNL2 expression (Fig. 2*C*), which suggested NMDAR-mediated reduction. We then examined MBNL2 expression under NMDA treatment and found that hippocampal neurons treated with 50 μM NMDA showed a similar pattern of MBNL2 downregulation as glutamate-treated neurons (Fig. 2*D*). Before NMDA stimulation, blockade of extracellular calcium influx by EGTA prevented NMDA-reduced MBNL2 expression (Fig. *2E*). The results suggest that NMDAR-mediated excitotoxicity induced MBNL2 degradation.

To further determine whether NMDA-reduced MBNL2 expression affects the developmental RNA processing transition, we analyzed the alternative splicing and polyadenylation of MBNL2 targets in glutamate and NMDA-treated neurons. The pattern of alternative splicing and polyadenylation in neurons with NMDAR-mediated excitotoxicity was dysregulated and similar to that in Mbnl2-depleted neurons (Fig. 1*E* and 2*F*), which suggests that NMDAR-mediated excitotoxicity reduced the MBNL2 expression and developmental pattern of MBNL2-regulated RNA processing. Therefore, we used NMDA to investigate the causal mechanism of NMDAR-reduced MBNL2 protein expression for the rest of the study.

### NMDAR-mediated calpain-2 activation causes MBNL2 degradation

To investigate how NMDAR-mediated excitotoxicity reduced MBNL2 protein level, we first determined whether MBNL2 downregulation was at the mRNA level. MBNL2 mRNA level was similar in neurons treated with glutamate or NMDA for 3 hr and unstimulated neurons (Fig. 3*A*), so NMDA-reduced MBNL2 expression was not due to downregulation at the mRNA level.

**Fig. 3.**
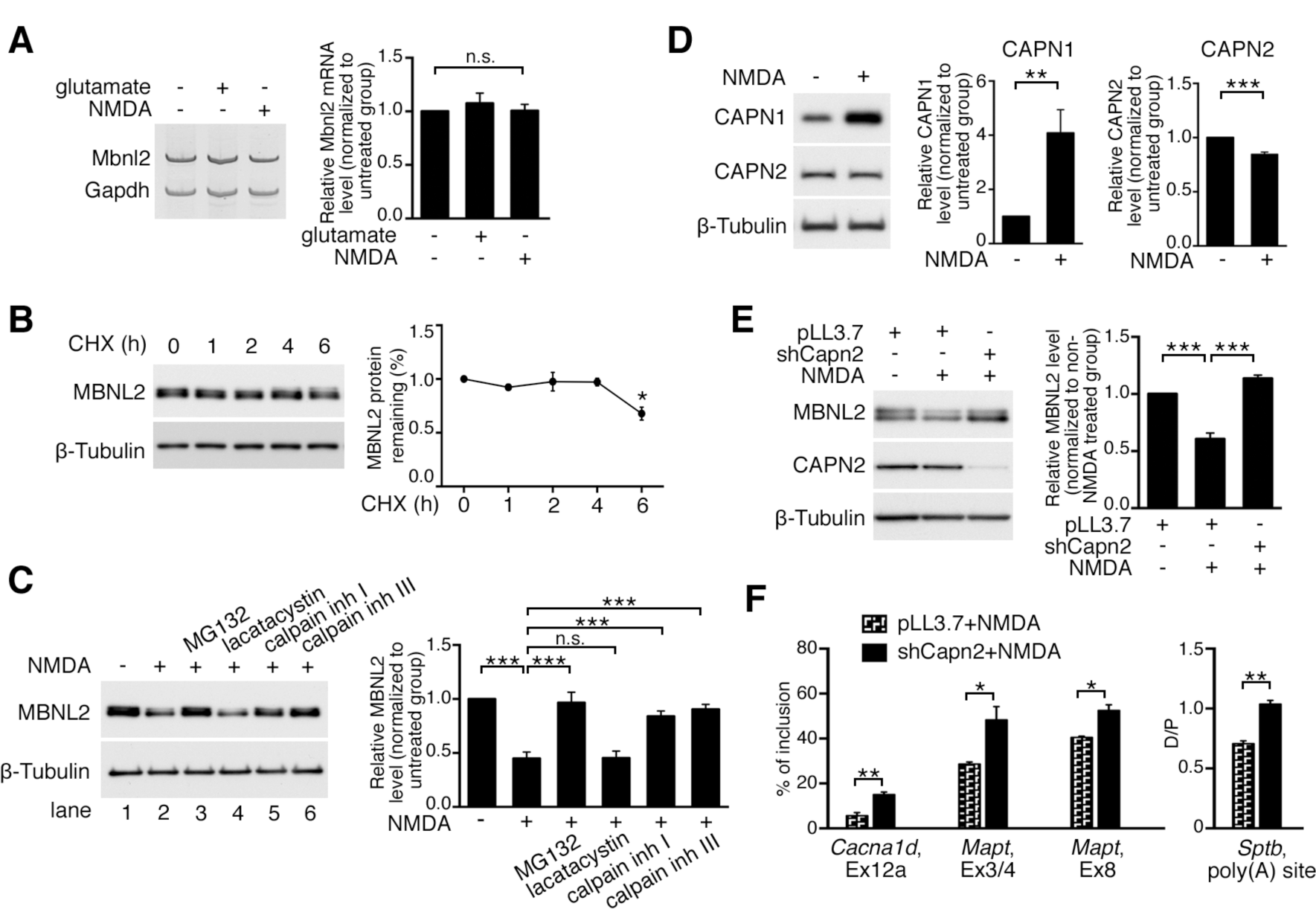
Calpain-2 activity is required for NMDAR-mediated MBNL2 degradation. (*A*) RT-PCR detection of Mbnl2 mRNA in neurons treated with glutamate or NMDA. Gapdh was a loading control. (*B*) Determination of MBNL2 protein stability with lysates from hippocampal neurons treated with cycloheximide (CHX) for the indicated times. (*C*) MBNL2 level detected by immunoblotting in neurons treated with combinations of drugs as indicated. (*D*) Change of CAPN1 and CAPN2 protein levels in NMDA-treated neurons. (*E*) CAPN2 knockdown preserved MBNL2 level in NMDA-treated neurons. Neurons were infected with lentivirus expressing CAPN2 shRNA (shCapn2) or control virus expressing pLL3.7. (F) Knockdown of CAPN2 preserved the developmental RNA processing transition in NMDA-treated neurons. Data are mean ± SEM of triplicate determinations. \**P*<0.05, \*\**P*<0.01,***P< 0.001, by one-way ANOVA (*A, B, C* and *E*) or student *t* test (*D* and *F*). n.s., not significant.

To determine whether reduced MBNL2 expression was regulated at the post-translational level, we first examined the protein stability of MBNL2 in mature neurons. In neurons treated with cycloheximide (CHX) to inhibit protein synthesis, MBNL2 protein was relatively stable and slightly reduced in level, ∼20%, after 6 hr of CHX treatment (Fig. 3*B*). We next determined whether reduced MBNL2 expression was due to proteasome degradation by pretreating neurons with two proteasome inhibitors, MG132 or lactacystin, before NMDA treatment. Pretreatment with MG132 but not lactacystin prevented the reduced MBNL2 expression (Fig. 3*C*, lanes 1-4). In addition to proteasome inhibition, MG132 also inhibits activities of other cysteine proteases including cathepsins and calpains (26). The lack of response to lactacystin pretreatment suggested that NMDA-reduced MBNL2 expression was not due to proteasome degradation. Calpains belong to a family of calcium-dependent cysteine proteases, and their activity can be induced by NMDAR-induced calcium influx (27, 28). We next assessed whether calpain was involved in regulating MBNL2 downregulation. We pretreated neurons with N-acetyl-L-leucyl-L-leucyl-L-norleucinal (calpain inhibitor I) or carbobenzoxy-valinyl-phenylalaninal (calpain inhibitor III) before NMDA treatment and found that both inhibitors prevented NMDA-reduced MBNL2 expression (Fig. 3*C*, lanes 5-6). Thus, NMDAR-mediated MBNL2 downregulation was likely via calpain activation.

To further determine whether NMDA-induced MBNL2 degradation was regulated by calpain, we tested whether calpain knockdown could prevent MBNL2 degradation under NMDA treatment. The major calpains expressed in neurons are calpain-1 and -2 (27). Both contain a distinct catalytic subunit, CAPN1 or CAPN2, respectively, and a common regulatory subunit, CAPNS1 (28). We first determined the levels of calpain-1 and -2 by examining those of CAPN1 and CAPN2 in response to NMDA treatment in cultured hippocampal neurons. In neurons treated with NMDA for 3 hr, total CAPN1 level was increased, whereas that of CAPN2 was slightly reduced as compared with that for unstimulated neurons (Fig. 3*D*). We next determined which calpain regulates MBNL2 degradation by depleting endogenous CAPN1 and CAPN2 by infection with a lentivirus expressing *Capn1*- or *Capn2*-specific shRNA, respectively (*SI Appendix*, Fig. S3A). In NMDA-treated neurons, CAPN1 knockdown failed to prevent MBNL2 degradation (*SI Appendix*, Fig. S3*B*), whereas CAPN2 knockdown prevented MBNL2 degradation after NMDA stimulation (Fig. 3*E*). We next determined whether depletion of CAPN2 could prevent NMDA-induced aberrant RNA processing regulated by MBNL2. In Capn2-depleted neurons treated with NMDA, the inclusion of *Cacna1d* exon 12a, *Mapt* exon 3/4 and exon 8 was increased (Fig. 3*F*) and the selection of *Sptb* alternative polyadenylation sites were toward the distal site, which indicated an inhibition of aberrant RNA processing on CAPN2 depletion in NMDA-treated neurons. These results suggest that calpain-2 regulated MBNL2 degradation and the developmental RNA processing transition.

### Nuclear translocation of calpain-2 results in nuclear MBNL2 degradation under neurodegenerative conditions

Calpain-2 is a cytosolic cysteine protease. Immunofluorescence staining and biochemical fractionation examination of CAPN2 localization in cultured hippocampal neurons showed a predominantly cytoplasmic distribution (*SI Appendix*, Fig. S4). However, MBNL2 distribution was relatively abundant in the nucleus of cultured hippocampal neurons (Fig. 2*B*, top). To determine how cytoplasmic-localized calpain-2 may mediate MBNL2 degradation under NMDA treatment, we wondered whether it was due to NMDA-induced nuclear translocation of calpain-2. We first examined the subcellular distribution of calpain-2 under NMDA treatment. The nucleus-to-cytoplasm ratio of CAPN2 immunoreactivity was increased in neurons treated with NMDA as compared with vehicle-treated control neurons (Fig. 4*A*). Using biochemical fractionation, we found that NMDA treatment increased CAPN2 level in the nuclear fraction but decreased it in the cytoplasmic fraction (Fig. 4*B*), which suggested a possible involvement of nuclear translocation. Although reduced MBNL2 expression was detected in both cytoplasmic and nuclear fractions, the extent of reduction was greater in the nuclear than cytoplasmic fraction (remainder: nuclear, 45%; cytoplasmic, 60%, Fig. 4*B*). We next examined the effect of CAPN2 knockdown on MBNL2 subcellular distribution under NMDA treatment. Knockdown of CAPN2 in NMDA-treated neurons preserved MBNL2 levels in both nuclear and cytoplasmic fractions (Fig. 4*C*), which suggests that calpain-2 undergoing nuclear translocation mediates MBNL2 degradation in both nucleus and cytoplasm.

**Fig. 4.**
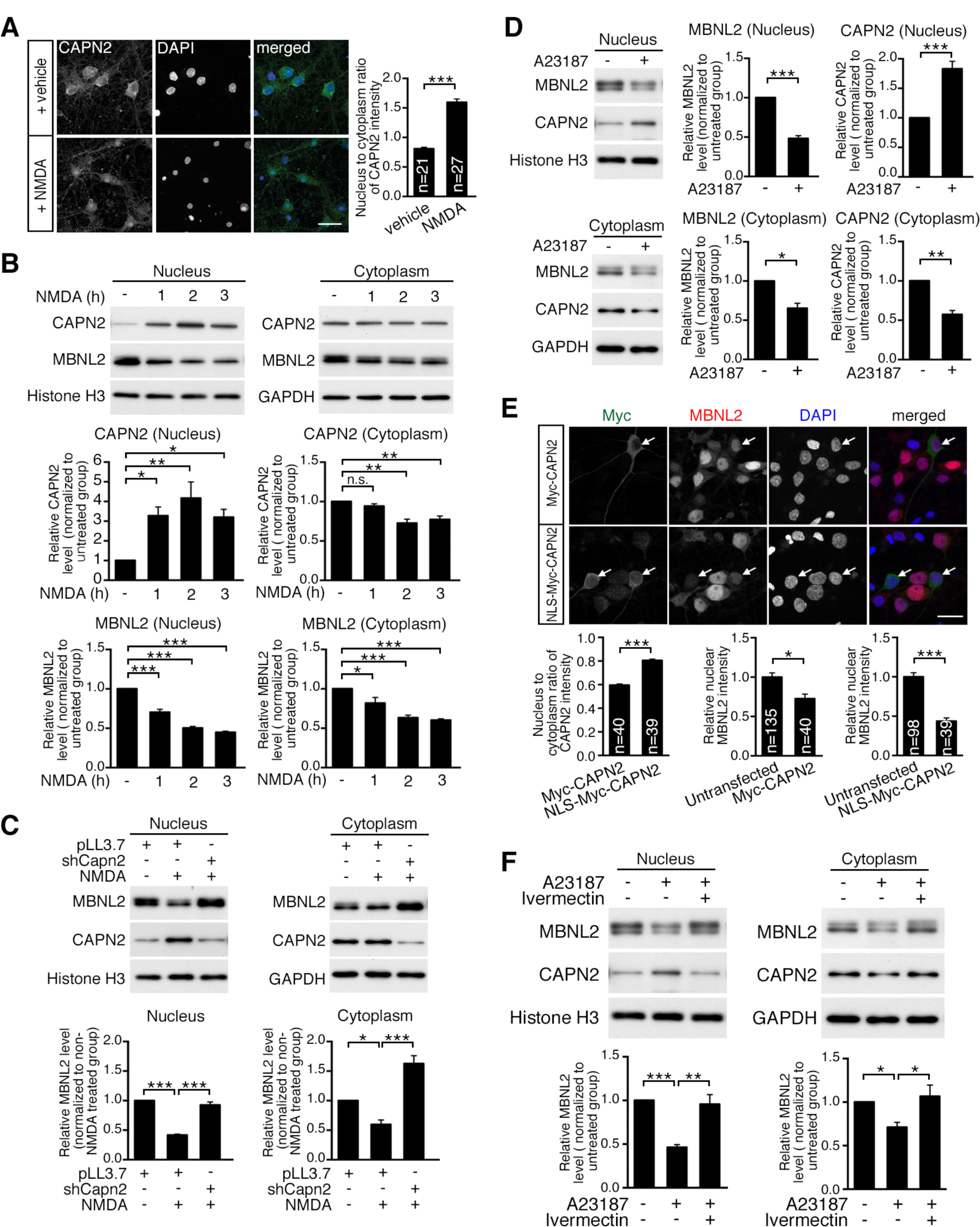
Nuclear translocation of calpain-2 results in nuclear MBNL2 degradation under neurodegenerative conditions. (*A*) NMDA treatment increased nuclear fraction of CAPN2. Quantification of nucleus-to-cytoplasm ratio of CAPN2 immunoreactivity at right. (Scale: 20 µm.) (*B*) Nuclear and cytoplasmic distribution of CAPN2 and MBNL2 in neurons treated with NMDA for the indicated times. Histone H3 and GAPDH were loading controls for nuclear and cytoplasmic fractions, respectively. Quantification of cytoplasmic and nuclear fractions of CAPN2 and MBNL2 relative to loading controls is indicated at the bottom. (*C*) CAPN2 knockdown preserved MBNL2 level in the nucleus and cytoplasm of NMDA-treated neurons. Neurons were infected with lentivirus expressing shCapn2 or pLL3.7 as control. (*D*) Detection of MBNL2 and CAPN2 level in the nuclear and cytoplasmic fractions of A23187-treated neurons. (*E*) Representative images of Myc and MBNL2 immunoreactivity in neurons transfected with Myc-CAPN2 (top) or NLS-Myc-CAPN2 (bottom). Arrows indicate neurons transfected with different constructs. Nucleus-to-cytoplasm ratio of Myc immunoreactivity in neurons transfected with Myc-CAPN2 and NLS-Myc-CAPN2 was quantified and shown at bottom. Nuclear MBNL2 level in neurons transfected with Myc-CAPN2 or NLS-Myc-CAPN2 were compared with that in untransfected neurons (at right). (Scale: 20 µm.) (*F*) Prevention of CAPN2 nuclear translocation by ivermectin preserved MBNL2 expression in A23187-treated neurons. Data are mean ± SEM from indicated n number of neurons (*A* and *E*) or 3 independent experiments (*B*-*D* and *F*). \**P*<0.05, \*\**P*<0.01, ***P< 0.001, by one-way ANOVA (*B, C* and *F*) or Student *t* test (*A, D*, and *E*). n.s., not significant.

In addition to NMDAR-mediated excitotoxicity, dysregulation of calcium homeostasis is implicated in the pathogenesis of neurodegeneration (29). We used a calcium ionophore, A23187, to treat neurons for mimicking the dysregulated calcium homeostasis to test whether aberrant calcium homeostasis may also induce CAPN2 nuclear translocation and MBNL2 reduction. Consistent with the findings seen in NMDA-treated neurons, in cultured neurons treated with A23187, MBNL2 expression was reduced in both cytoplasmic and nuclear fractions associated with an increase in CAPN2 level in the nuclear fraction and a decrease in CAPN2 level in the cytoplasmic fraction (Fig. 4*D*). In addition, knockdown of CAPN2 in neurons treated with A23187 preserved MBNL2 level (*SI Appendix*, Fig. S5A) and prevented the reversal of alternative splicing of *Cacna1d* exon 12a, *Mapt* exon 3/4 and exon 8 to the developmental pattern (*SI Appendix*, Fig. S5B).

We next investigated whether the nuclear localization of CAPN2 was involved in the reduced MBNL2 nuclear fraction. We generated an Myc-tagged CAPN2 mutant construct (NLS-Myc-CAPN2) with addition of a nuclear localization signal at the N-terminus of CAPN2 to increase the nuclear fraction of CAPN2. The wild-type Myc-tagged CAPN2 (Myc-CAPN2) mainly localized in the cytoplasm. Also, the expression of Myc-CAPN2 in cultured hippocampal neurons reduced endogenous MBNL2 level as compared with in untransfected neurons (Fig. 4*E*, ∼27%). Expression of the mutant CAPN2 increased the nuclear fraction of CAPN2 as examined by the nucleus-to-cytoplasm ratio of Myc immunoreactivity and further decreased the level of endogenous MBNL2 (Fig. 4*E*, ∼56%).

Next, we determined whether blocking CAPN2 nuclear translocation by inhibiting importin-mediated nuclear import with ivermectin (30) would prevent MBNL2 degradation. Pretreatment with ivermectin prevented A23187-reduced MBNL2 expression and CAPN2 nuclear translocation (Fig. 4F), that suggests a requirement of nuclear localization of CAPN2 for MBNL2 degradation in the nucleus. Therefore, our results suggest that neurodegenerative conditions caused CAPN2 nuclear translocation associated with reduced MBNL2 expression and re-induction of a developmental RNA processing program.

### Expression of calpain-2 follows a nuclear-to-cytoplasm transition during neuronal maturation

The reduced MBNL2 level associated with reversal of developmental RNA processing suggested re-induction of a developmental program under neurodegenerative conditions (Fig. 1*E*). We wondered whether calpain-2 nuclear translocation under neurodegeneration also represents a reversal of the developmental transition associated with MBNL2 expression. To address this question, we examined the cellular distribution of CAPN2 in cultured hippocampal neurons at an early and mature stage. CAPN2 was distributed in both the cytoplasm and nucleus of cultured neurons at 2 DIV but was mainly distributed in the cytoplasm at 16 DIV (Fig. 5A). As compared with CAPN2 distribution in neurons at 2 DIV, the ratio of nucleus-to-cytoplasm fraction of CAPN2 immunoreactivity was reduced in neurons at 16 DIV, which suggests a developmental transition from nucleus-to-cytoplasm distribution during neuronal maturation. Our results suggest that the neurodegenerative condition likely induced a developmental transition for CAPN2 from the cytoplasm to nucleus, thus resulting in MBNL2 reduction accompanied by reversal of the developmental RNA processing transition.

**Fig. 5.**
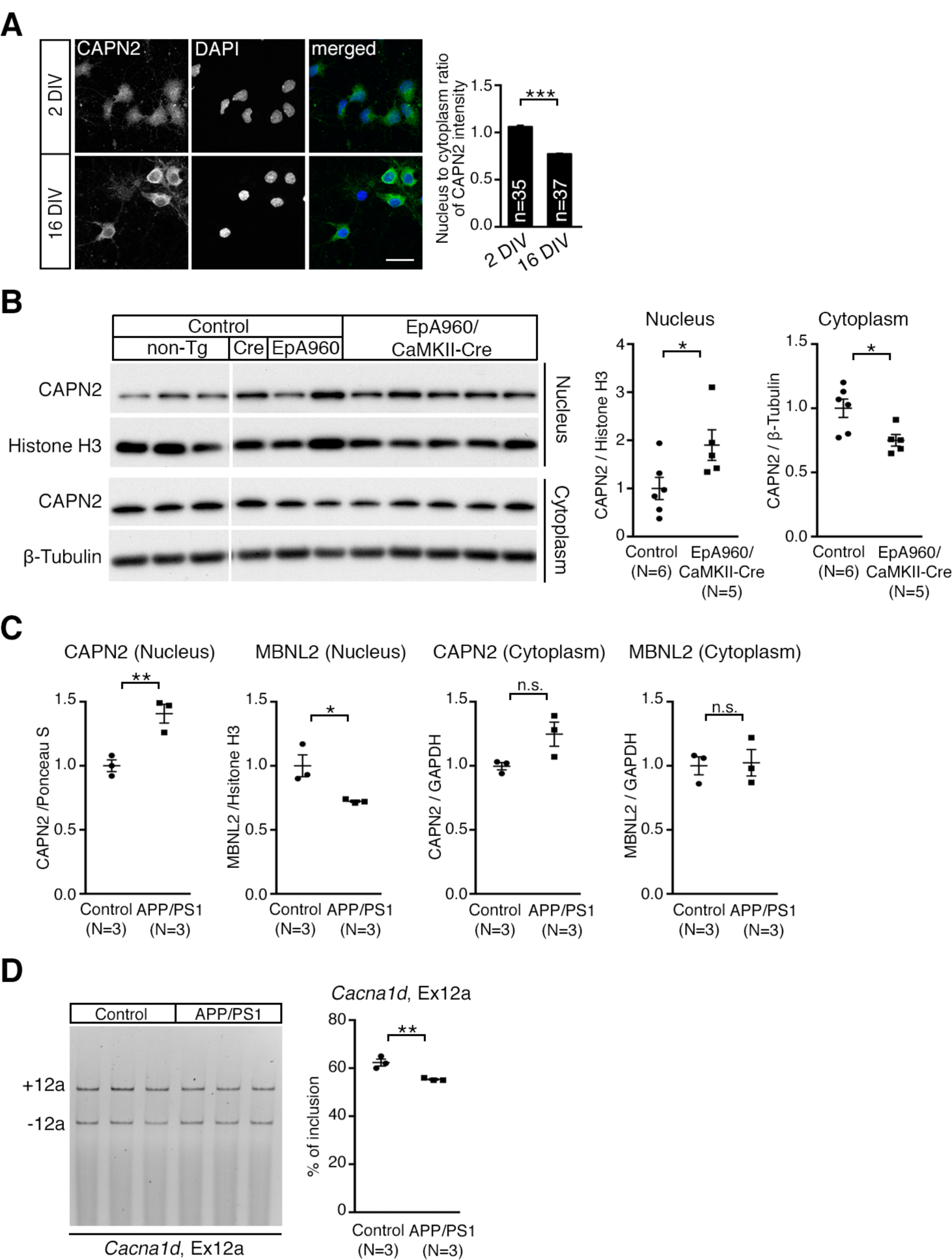
Increased nuclear CAPN2 associated with reduced MBNL2 level and re-induction of developmental RNA processing program in neurodegeneration. (A) Representative images of CAPN2 immunoreactivity in neurons at 3 and 16 DIV. Quantification of nucleus-to-cytoplasm ratio of CAPN2 immunoreactivity at right. (Scale: 20 µm.) (B) CAPN2 content in nuclear and cytoplasmic fractions of lysates from control and EpA960/CaMKII-Cre brains. Brains from animals with different genotypes including non-Tg, CaMKII-Cre (Cre) and EpA960 brains were used as controls. (C) Detection of CAPN2 and MBNL2 level in nuclear and cytoplasmic fractions of cortical lysates from non-Tg (Control) and APP/PS1 mice at age 10 months. (D) Alternative splicing of *Cacna1d* exon 12a in non-Tg and APP/PS1 cortical tissues. Data are mean ± SEM of indicated number of neurons (n) or animals (N). \**P*<0.05,\*\**P*<0.01, ***P< 0.001, by Student *t* test. n.s., not significant.

### Re-induced developmental program of CAPN2 and MBNL2 expression patterns and MBNL2-regulated RNA processing in mouse brains with neurodegeneration

A brain-specific mouse model for DM1, EpA960/CaMKII-Cre, exhibited MBNL2 downregulation in both cytoplasm and nucleus (20). Reduced MBNL2 expression accompanied by aberrant alternative splicing occurred after axon and dendrite degeneration. We wondered whether nuclear translocation of CAPN2 also occurred in the EpA960/CaMKII-Cre brain. The cytoplasmic fraction of CAPN2 level was decreased and the nuclear fraction was increased in the EpA960/CaMKII-Cre brain (Fig. 5*B*), whereas the subcellular distribution of CAPN1 was unchanged among all animals (*SI Appendix*, Fig. S6). The results showed an association of calpain-2 nuclear translocation with reduced MBNL2 level in the EpA960/CaMKII-Cre brain, a similar pattern to that of neurons with neurodegenerative features including NMDAR-mediated excitotoxicity and calcium homeostasis dysregulation.

We next wondered whether the CAPN2 nuclear translocation associated with reduced MBNL2 level also occurred in the APP/PS1 brain, a mouse model for AD (31). In the APP/PS1 brain, the formation of amyloid plaques was followed by deficits in learning and memory, synaptic dysfunction and neuronal loss. In 10-month-old APP/PS1 brains, the nuclear fraction showed increased CAPN2 level associated with reduced MBNL2 expression as compared with control brains, with no change in cytoplasmic fraction of CAPN2 and MBNL2 (Fig. 5*C*). We further examined MBNL2-regulated alternative splicing and found decreased inclusion of *Cacna1d* exon 12a in the APP/PS1 brain, showing a reversal of developmental transition pattern (Fig. 5*D*). These results suggest that the increased nuclear calpain-2 level accompanied by reduced MBNL2 level and re-induction of the developmental RNA processing program is likely a common feature of neurodegeneration.

## Discussion

The MBNL protein family plays an important role in promoting differentiation by regulating the transition of developmental RNA processing (14, 16) and is involved in the pathogenesis of DM1 (32). Investigations with mouse models expressing expanded CUG RNA or depletion of endogenous MBNL proteins have revealed the distinct roles of MBNL1 and MBNL2 in contributing to DM1 features (14, 17, 20). In a brain-specific DM1 mouse model, EpA960/CaMKII-Cre, degeneration was followed by reduced MBNL2 expression accompanied by aberrant alternative splicing. The correlation between neurodegeneration and reduced MBNL2 expression has not been determined. Whether the dysregulated transition of developmental RNA processing could be a common feature of neurodegeneration has not been tested. In the present study, we found the expression and function of MBNL2 associated with progression of neuronal maturation. Depletion of MBNL2 in cultured hippocampal neurons impaired dendrite morphogenesis and caused dysregulated transition of developmental RNA processing, which suggests the requirement for MBNL2 in promoting differentiation. We also showed that neurodegenerative conditions including NMDAR-mediated excitotoxicity and dysregulated calcium homeostasis induced MBNL2 downregulation at the post-translational level in cultured hippocampal neurons. Reduced MBNL2 protein level was mediated by not proteasome degradation but rather calcium-dependent calpain activity. Comparable to the effects induced by expanded CUG RNA, NMDA-reduced MBNL2 expression resulted in reversal of MBNL2-regulated alternative splicing and polyadenylation to the developmental pattern. Depletion of endogenous CAPN2 prevented NMDA- or A23187-induced MBNL2 degradation and aberrant RNA processing. Neurodegenerative conditions induced nuclear translocation of cytoplasmic calpain-2, thereby degrading the nuclear fraction of MBNL2. Moreover, we showed that distribution of calpain-2 was developmentally regulated and exhibited a nuclear-to-cytoplasm transition during neuronal maturation. The nuclear translocation of calpain-2 induced by neurodegenerative stimulation resembled its developmental transition. Combined with our present study, the EpA960/CaMKII-Cre and APP/PS1 mouse models for DM1 and AD neurodegeneration, respectively, showed an increase of nuclear calpain-2 associated with reduced MBNL2 level and dysregulated alternative splicing. Thus, our findings provide a novel mechanism for calpain-mediated MBNL2 degradation in the neural pathogenesis of DM1. It also suggests that the neurodegenerative condition may re-induce a developmentally regulated program in which calpain-mediated MBNL2 degradation associated with aberrant RNA processing may represent a common pathogenic pathway in neurodegenerative disorders.

The major molecular feature of DM1 is reversion of alternative splicing and polyadenylation to the embryonic pattern (13, 14, 33). Indeed, re-induction of fetal/neonatal gene program has been reported under several pathological conditions. After myocardial infarction CELF1 is upregulated, which is normally down-regulated in adult heart, resulting in mRNA degradation of genes involved in cardiac conduction and contractility (34). Upon toxin-induced injury, adult hepatocyte regeneration is associated with reactivation of neonatal translation and splicing programs (35). Similarly, after spinal cord injury, transcriptional profile of corticospinal tract (CST) motor neurons temporarily exhibits an embryonic pattern (36). Grafting of spinal-cord-derived neural progenitor cells sustains the embryonic transcriptional profile in CST motor neurons enabling the regeneration of corticospinal axons (36). Aerobic glycolysis or Warburg effect in cancer cells is a reprogrammed metabolism resulting from increased glucose uptake and lactate production in the presence of oxygen (37, 38). Expression of embryonic isoform of pyruvate kinase PKM2 is required for aerobic glycolysis providing the advantage for tumor growth (37, 38). Reversion of pyruvate kinase to adult isoform PKM1 decreases lactate production and increases oxygen consumption resulting in tumor growth reduction (37, 38). In the present study, we showed that nuclear localization of calpain-2 and low expression level of MBNL2 seen in neurodegenerative conditions recapitulate their developmental expression pattern. Therefore, re-induction of fetal gene program is likely a common feature induced by the pathological condition. Whether re-induction of fetal gene program in adult tissues provides a beneficial effect or contributes to adverse phenotypes is worthy of investigation.

MBNL2 has been found involved in regulating splicing switches during postnatal neuronal maturation (39). Consistently, we showed that the temporal expression pattern of MBNL2 in the dentate gyrus of the hippocampal region paralleled the neuronal differentiation pattern, with MBNL2 initially expressed in the outer layer of the dentate gyrus and increasingly following the ectal-to-endal pattern during postnatal development. The reduced expression of MBNL2 in the DM1 brain and neurons subjected to excitotoxicity or dysregulated calcium homeostasis was reminiscent of its low expression during early developmental stages. Thus, changes in the functions of regulating RNA processing and the expression of MBNL2 under excitotoxic conditions are related. Previously, we showed that MBNL1 was predominantly distributed in the cytoplasm in the embryonic mouse brain and became localized in the nucleus and cytoplasm in the adult brain (20). The predominant cytoplasmic localization of MBNL1 in the embryonic brain and increased nuclear distribution of both MBNL1 and MBNL2 in the postnatal brain suggest that other RBPs may be involved in regulating embryonic RNA processing events and are worthy of investigation.

Our finding of aberrant MBNL2-regulated RNA processing in cultured neurons caused by NMDAR-activated calpain was consistent with that in brains of individuals with DM1 and MBNL2-knockout animals, which suggests a neuron-specific regulatory role for MBNL2. In the DM1 brain, decreased inclusion of *Mapt* exon 10, encoding the R2 of microtubule-binding repeat domain, increased the expression of the fetal tau isoform 3R and reduced that of the adult tau isoform 4R (8, 9). Tau protein regulates the assembly and stability of microtubules (40, 41). 3R tau has less activity than 4R tau in binding to tubulin/microtubules (42), which results in an increase in the unbound form of 3R tau, a favorable substrate for phosphorylation (43, 44). Hyperphosphorylation of tau leads to the formation of neurofibrillary tangles, a critical pathogenic event of dementia (45). Of note, increased tau 3R:4R protein ratio was reported in Pick disease (46) and Down syndrome (43), which suggests a potential regulatory mechanism by MBNL2-regulated alternative splicing in other neurodegenerative disorders.

Calpain activation has been identified in several neurodegenerative disorders, and most calpain substrates are in the cytoplasm (27, 47). We showed that MBNL2 is likely a substrate of calpain-2 in the nucleus in the context of neurodegeneration, and nuclear translocation of calpain-2 was required for its function to degrade MBNL2. Further study of how calpain-2 nuclear translocation is regulated under the excitotoxic condition is needed. We showed that NMDAR-activated excitotoxicity induced the levels of calpain-1 and -2. However, genetic knockdown of only calpain-2 but not calpain-1 resulted in reversal of MBNL2 degradation and aberrant MBNL2-regulated RNA processing. Calpain-1 and -2 may regulate different substrates because of differences in their structures and dependence on calcium concentration for activation (28, 48), which suggests the distinct role of different calpains in the progression of neurodegenerative disorders. Our results of calpain-2-mediated MBNL2 degradation and aberrant RNA processing may suggest a converging pathway in the pathogenesis of calpain-related neurodegenerative disorders.

Neurodegenerative conditions including NMDAR-mediated excitotoxicity and dysregulated calcium homeostasis activate calpain-2 and trigger its nuclear translocation in neurons. In the EpA960/CaMKII-Cre brain, calpain-2 level was increased in the nucleus but decreased in the cytoplasm, which is consistent with the pattern of calpain-2 nuclear translocation in neurons under calcium-dependent neurodegenerative conditions. These results suggest the involvement of calcium-dependent neurotoxicity and calpain-2 activation in the brain pathogenesis of DM1. A previous finding demonstrates that expression of GLT1 glutamate transporter is reduced in astrocytes, disrupting the glutamate uptake by astrocytes and causing glutamate neurotoxicity on the Purkinje cells in the cerebellum of DMSXL mice, a DM1 model carrying a human *DMPK* transgene containing more than a thousand of CTG repeats (49). This finding also implicates the role of glutamate-mediated excitotoxicity in the pathogenesis of DM1. Together, investigation of dysregulation of glutamate concentration and calcium homeostasis in DM1 brain is needed and may provide important insights for developing therapeutic approaches for DM1 brain pathology.

## Materials and Methods

### Animals and tissues

The EpA960/CaMKII-Cre mice were described previously (20). F1 offspring from EpA960 mice crossed with CaMKII-Cre mice were used in experiments. The APP/PS1 mice were originally from the Jackson Laboratory [B6;C3-Tg (APPswe, PSEN1dE9) 85Dbo/Mmjax]. Male APP/PS1 mice and age-matched non-Tg control littermates at 10 months of age were used in this study. All animals were maintained on a standard 12-h light/dark cycle with light on at 8 AM. Food and water were available *ad libitum*. All animal experiments were performed with the approval of the Academia Sinica Institutional Animal Care and Utilization Committee in strict accordance with its guidelines and those of the Council of Agriculture Guidebook for the Care and Use of Laboratory Animals.

### Primary hippocampal neuron culture, drug treatment, virus infection and transfection

Hippocampal neurons were cultured from rat embryos at embryonic day 18-19 as described (21). In total, 200,000 neurons per well were plated in 12-well plates containing coverslips coated with poly-L-lysine (1 mg/ml). Neurons at 19 days of *in vitro* culture (DIV) were pretreated with tetrodotoxin (TTX, 1 µM) for 12-16 hr to reduce endogenous synaptic activity. Cells were stimulated with glutamate (20 µM), NMDA (50 µM) or A23187 (4 µM) at 20 DIV for 1-3 hr. 6,7-dinitro-2,3-dihydroxyquinoxaline (DNQX) (10 µM), AP5 (50 µM), MG132 (10 µM), lactacystin (10 µM), calpain inhibitor I (40 µM), calpain inhibitor III (40 µM), EGTA (2 mM) or ivermectin (1 µM) was added 30 min before stimulation for 3 hr or the indicated times. To study the effect of blocking protein synthesis, cells were incubated with cycloheximide (CHX; 10 µM). After drug treatment, neurons were lysed for immunoblotting or fixed for immunocytochemistry.

For genetic-knockdown experiments, hippocampal neurons at 10 DIV were infected with lentivirus expressing target-specific short hairpin RNA (shRNA) for 24 hr and harvested at 16 DIV for protein or RNA extraction. To study the knockdown effect on NMDA stimulation, virus-infected cells at 16 DIV were pre-treated with TTX for 12-16 hr before NMDA, then lysed for protein or RNA extraction.

Transfection with calcium phosphate precipitation was performed at 11 DIV, followed by fixation at 14 DIV and immunofluorescence staining. All transfection experiments and drug treatments with hippocampal neurons were repeated 3 times on different batches of neuronal preparations from embryos derived from different pregnant rats.

### Subcellular fractionation and immunoblotting

Subcellular fractionation for cytoplasmic and nuclear protein from cultured hippocampal neurons was performed as described (34). Briefly, cells were lysed on ice by using gentle lysis buffer (10 mM HEPES, pH 7.5, 3 mM MgCl_2_, 10 mM NaCl and 0.5% NP-40) and centrifuged at 2,000 *g* for 8 min. The supernatants were collected as the cytoplasmic fraction and pellets were suspended by using gentle lysis buffer containing 0.1% SDS. The suspensions were sonicated for up to 3.5 min and centrifuged at 12,000 rpm for 20 min. The supernatants were harvested as the nuclear fraction.

For total protein extraction from cultured hippocampal neurons, cells were lysed by using SDS-sample buffer (188 mM Tris, pH 6.8, 3% SDS, 3% glycerol and 0.01% bromophenoblue). The lysates were incubated at 55°C for 10 min and resolved by western blot analysis. For immunoblotting analysis of mouse brains, tissues were homogenized by tight douncing in buffered sucrose (4 mM HEPES, 320 mM sucrose, 2 mM DTT, 2 mM MgCl_2_, 1 mM EDTA and protease inhibitors), lysed in buffered sucrose containing 1% SDS, then sonicated and centrifuged at 13,000 rpm for 10 min; the supernatant was collected as total protein lysates for immunoblotting. The subcellular fractionation of mouse brains was previously described (20); tissues were homogenized by loose douncing in buffered sucrose and centrifuged at 800 *g* for 10 min. The supernatants were collected as the cytoplasmic fraction and pellets were re-suspended by using buffered sucrose containing 1% SDS followed by sonication and collection as the nuclear fraction.

### RNA preparation, RT-PCR and splicing analysis

Total RNA was extracted by using TRIzol reagent (Invitrogen) and used for cDNA synthesis followed by PCR as described (50). Primers used for MBNL2 and GAPDH amplification were Mbnl2-F, 5’-CTCTGCAGCAACAACTCCTGC-3’; Mbnl2-R, 5’-TAGCAGAACTAG CCTTAGGG-3’; Gapdh-F, 5’-TGCACCACCAACTGCTTA-3’; Gapdh –R, 5’-TAGCAGAA CTAGCCTTAGGG-3’. For RT-PCR splicing analysis, the primers used for alternative splicing of *Cacna1d* exon 12a, *Mapt* exons 3 and 8 and alternative polyadenylation of *sptb* were previously described (33). PCR products were analyzed with 5% nondenaturing polyacrylamide gels. Quantification of the percentage of exon inclusion was as described (51).

### Immunofluorescence staining and immunohistochemistry

For the detection of endogenous protein expression, immunofluorescence staining was performed as described (21). Briefly, cells were fixed with phosphate buffered saline (PBS) containing 4% paraformaldehyde (PFA) and 4% sucrose for 15 min at room temperature. Fixed cells underwent permeabilization and blocking in 3% normal goat serum (NGS) in tris-buffered saline (TBS) containing 0.2% Triton X-100 for 1 hr, followed by incubation with primary antibodies overnight in incubation solution (1% NGS and 0.2% Triton X-100 in TBS), then with secondary antibodies conjugated with Alexa Fluor (Invitrogen) and with DAPI for nuclear labeling for 2 hr. After washing, cells were mounted with Fluoro-gel (EMS) for imaging.

Immunohistochemistry of mouse brain sections was as described (20). Briefly, mice were anesthetized and transcardially perfused with PBS and 4% PFA sequentially. Brain tissues were post-fixed in 4% PFA for 16 hr at 4°C and sectioned by vibratome sectioning. To detect endogenous MBNL2 expression, brain sections first underwent bleaching with 3% H_2_O_2_ in TBS for 10 min and permeabilization in TBS containing 0.5% Triton X-100 for 10 min. After incubation with blocking solution (3% NGS and 0.2% Triton X-100 in TBS) for 1 hr, sections were incubated with anti-MBNL2 antibody in incubation solution (1% NGS and 0.2% Triton X-100 in TBS) overnight at 4°C. After washing, sections were incubated with biotinylated secondary antibody (Vector Laboratories) for 2 hr, then with ABC reagent (Vector Laboratories) for 2 hr, followed by signal development by using the SG substrate kit (Vector Laboratories). Sections were counterstained with Nuclear Fast Red to label nucleus and coverslipped by using permount mounting medium (Vector Laboratories).

Fluorescent images were acquired under a fluorescent microscope (AxioImager M2, Carl Zeiss) equipped with a 40x/0.75 objective lens (Plan-Apochromat, Carl Zeiss) or a confocal laser scanning microscope (LSM700, Carl Zeiss) equipped with a 40x/1.4/oil objective lens (Plan-Apochromat, Carl Zeiss). Images for immunohistochemistry were acquired under an upright microscope (BX51, Olympus) equipped with 10x/0.4 or 20x/0.75 objective lenses (UPlanSApo, Olympus).

For all experiments with cultured hippocampal neurons, the image acquisition involved the same settings for experimental groups, with transfections performed at the same time and used for comparison.

### Antibodies

Antibodies used were anti-MBNL2 (sc-136167, 1:2500, for immunoblotting and 1:400 for immunostaining, Santa Cruz Biotechnology), anti-calpain-1 (CAPN1; ab39170, 1:5000, Abcam), anti-CAPN2 (sc-373966, 1:10000 for immunoblotting and 1:250 for immnostaining, Santa Cruz Biotechnology), anti-NeuN (ab177487, 1:1000 for immunostaining, Abcam), anti-β-tubulin (NB600-936, 1:20000, Novus), anti-GAPDH (MAB374, 1:40000, Millipore), anti-Histone H3 (sc-8654, 1:5000, Santa Cruz Biotechnology).

### Plasmids

The target-specific shRNA clones, TRCN 0000072224 (shLacZ, CGCGATCGTAATCACC CGAGT), TRCN0000178919 (shMbnl2-1, CAACACCGTAACCGTTTGTAT), TRCN0000102459 (shMbnl2-2, CAAAGAGACAAGCACTTGAAA), TRCN0000030665 (shCapn1, GCCGTGGACTTTGACAACTTT) were purchased from an RNAi core facility (Academia Sinica). The Capn2-specific shRNA clone (shCapn2, GGAGTGCCCTTCTGG AGAA) and the construct of rat CAPN2 sequence in pcDNA3.1 (52) was kindly provided by Dr. Yi-Shuian Huang (Academia Sinica). The rat CAPN2 sequence was sub-cloned into using the forward primer, AATTGGTACCTCATGGCGGGCATCGCGATG, and the reverse primer, AATGTCGACTCAGAGTACTGAAAAACTC, into GW1-Myc backbone (Myc-CAPN2). The nuclear localization signal (NLS) of SV40 large-T antigen was inserted using sense primer, GCTTGCCACCATGCCTAAGAAGAAGCGCAAGGTTGAGCAAAAGCTG, and antisense primer, CAGCTTTTGCTCAACCTTGCGCTTCTTCTTAGGCATGGTGGCA AGC, at the N-terminus of Myc tagged-CAPN2 (NLS-Myc-CAPN2).

### Statistical analysis

Data are expressed as mean ± SEM. Statistical analysis was performed with SigmaPlot 12.5 (Systat Software Inc.). Unpaired two-tail Student *t*-test was used for comparing two groups, and one-way ANOVA followed by Tukey Test for comparing multiple groups. *P*<0.05 was considered statistically significant.

## Acknowledgments

We thank Dr. Yi-Shuian Huang (Institute of Biomedical Sciences, Academia Sinica) for sharing reagents. This work was supported by the Institute of Biomedical Sciences, Academia Sinica [to G.-S.W]. Funding for open access charge: [Institute of Biomedical Sciences, Academia Sinica].

## Author Contributions

LHW and GSW conceived the project, designed and performed the experiments and wrote the paper. YML and CYL gathered the tissues from animal models. YC advised on the project and manuscript.

