## Supplementary information for "Neurodegeneration induces a developmental RNA processing program by calpain-mediated MBNL2 degradation"

**Authors/Affiliations:**

Lee-Hsin Wang[a,b], Yu-Mei Lin[b], Chien-Yu Lin[b], Yijuang Chern[a,b,c] and Guey-Shin Wang[a,b,c,*]

[a]Taiwan International Graduate Program in Interdisciplinary Neuroscience, National Yang-Ming University and Academia Sinica, Taipei, Taiwan

[b]Institute of Biomedical Sciences, Academia Sinica, Taipei 11529, Taiwan

[c]Program in Molecular Medicine, National Yang-Ming University and Academia Sinica, Taipei, Taiwan

***To whom correspondence should be addressed:**

Guey-Shin Wang, Ph.D

Institute of Biomedical Sciences,

Academia Sinica

128, Section 2, Academia Rd.

Taipei 11529, Taiwan.

**Supplementary Information**

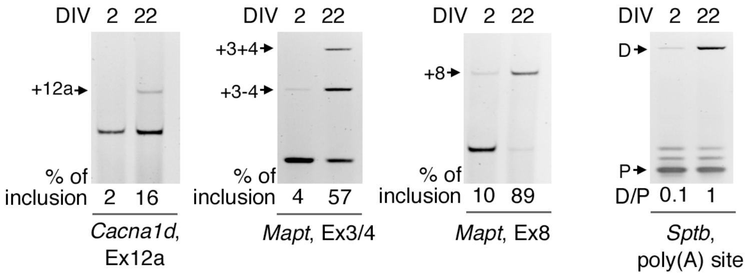


Figure S1

**Fig. S1.** Developmental changes of MBNL2-regulated RNA processing in hippocampal neurons. The alternative splicing of *Cacna1d* exon 12a, *Mapt* exon3/4 and exon8, and the alternative polyadenylation of *Sptb* as MBNL2 targets were examined in hippocampal neurons at 2 and 22 DIV. Quantification of the inclusion of *Cacna1d* exon 12a, *Mapt* exons 3/4 and 8, and the distal (D) to proximal (P) polyA utilization of *Sptb*. Data are mean of 2 independent experiments.

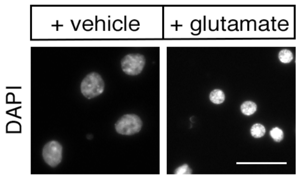


Figure S2

**Fig. S2.** Characterization of the effect of glutamate treatment on the condensation of nucleus in matured hippocampal neurons. DAPI was used for nuclear staining. Scale: 20 μm.

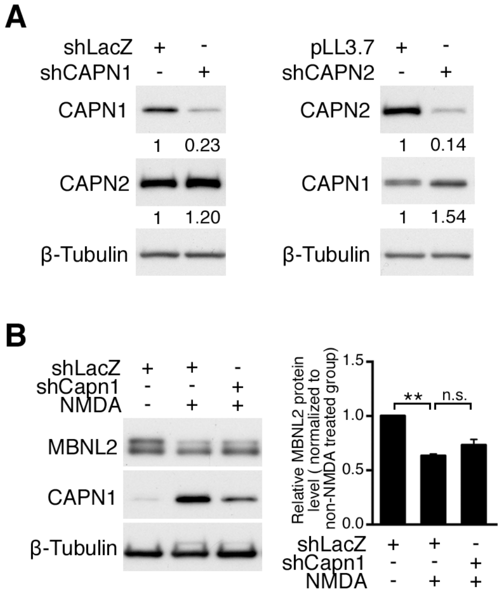


Figure S3

**Fig. S3.** (A) The efficiency and specificity of CAPN1 and CAPN2 knockdown in cultured hippocampal neurons. Lentivirus expressing specific shRNAs to knock down CAPN1 (shCapn1) or CAPN2 (shCapn2) was introduced into cultured hippocampal neurons. Virus that expressed shLacZ or pLL3.7 was used as a control. Levels of CAPN1 and CAPN2 were determined by immunoblotting analysis. β-tubulin was used as a loading control. Relative CAPN1 and CAPN2 level normalized to β-tubulin is shown as indicated numbers. (B) Effect of CAPN1 knockdown on MBNL2 protein expression in neurons treated with or without NMDA for 3 hr. Neurons were infected with lentivirus expressing CAPN1-specific shRNA (shCapn1) or control virus expression shLacZ. Relative MBNL2 levels normalized to β-tubulin are indicated. Data are mean ± SEM of triplicate determinations. **$P<0.01$, by one-way ANOVA. n.s., not significant.

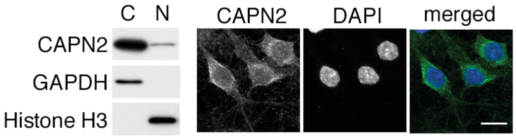


Figure S4

**Fig. S4.** Subcellular distribution of endogenous CAPN2 in mature hippocampal neurons by biochemical fractionation (left) and immunofluorescence staining (right). DAPI was used for nuclear staining.

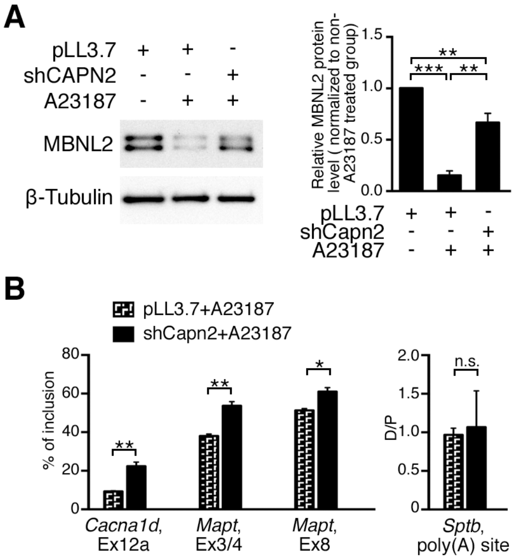


Figure S5

**Fig. S5.** Effects of CAPN2 knockdown on MBNL2 protein level (A) and MBNL2-regulated targets (B) in neurons treated with A23187. Lentivirus expressing shCapn2 shRNA to knock down CAPN2 (shCapn2) or control pLL3.7 shRNA was introduced into cultured hippocampal neurons. Data are mean ± SEM of triplicate determinations. **P*<0.05, ***P*<0.01, ****P*<0.001, by one-way ANOVA.

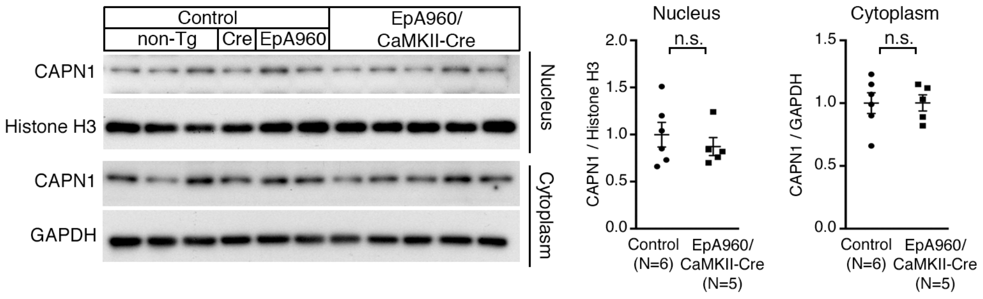


Figure S6

**Fig. S6.** CAPN1 expression pattern in the EpA960/CaMKII-Cre brains. Detection of CAPN1 in nuclear and cytoplasmic fractions of lysates from control and EpA960/CaMKII-Cre brains. Brains from animals with different genotypes including non-Tg, CaMKII-Cre (Cre) and EpA960 brains were used as controls. Data are mean ± SEM of indicated number of neurons animals (N). Statistical analysis is done by using Student *t* test. n.s., not significant.

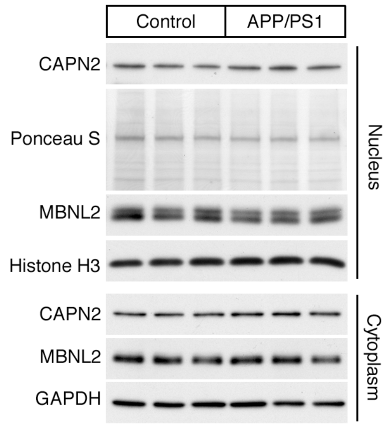


Figure S7

**Fig. S7.** Detection of CAPN2 and MBNL2 in nuclear and cytoplasmic fractions of lysates from control and EpA960/CaMKII-Cre brains. Non-Tg mice were used as control. Ponceau S of the same membrane for detecting nuclear CAPN2 was a loading control for CAPN2 in the nucleus. Histone H3 and GAPDH were loading controls for nuclear and cytoplasmic fractions, respectively.
